# Early alterations of large-scale brain networks temporal dynamics in young children with autism

**DOI:** 10.1101/2020.09.24.311423

**Authors:** Aurélie Bochet, Holger Franz Sperdin, Tonia Anahi Rihs, Nada Kojovic, Martina Franchini, Reem Kais Jan, Christoph Martin Michel, Marie Schaer

## Abstract

Disruption of large-scale brain networks is associated with autism spectrum disorders (ASD). Recently, we found that directed functional connectivity alterations of social brain networks are a core component of atypical brain development at early developmental stages in ASD (Sperdin et al., 2018). Here, we investigated the spatio-temporal dynamics of whole-brain neuronal networks at a subsecond scale in 90 toddlers and preschoolers (47 with ASD) using an EEG microstate approach. Results revealed the presence of five microstate classes that best described the entire dataset (labeled as microstate classes A-E). Microstate class C related to the Default Mode Network (DMN) occurred less in children with ASD. Analysis of brain-behavioural relationships within the ASD group suggested that a compensatory mechanism from microstate C was associated with less severe symptoms and better adaptive skills. These results demonstrate that the temporal properties of some specific EEG microstates are altered in ASD at early developmental stages.

## INTRODUCTION

High-density electroencephalography (EEG) represents a powerful mean to explore the brain’s physiological activity at a large-scale level in pediatric population (Michel & Murray, 2012). Recording the brain activity during a task or at rest in very young children with autism spectrum disorders (ASD) is challenging. However, early identification of brain alterations is important as it provides insights onto the brain mechanisms that lead to their clinical behavioral phenotype. Ultimately, increasing our understanding on how differently the brain develops during childhood years can help clinicians to adapt and use more tailored therapies early in life when the brain is most plastic and thus responsive to behavioral treatment.

Recently, by combining high density EEG and eye-tracking, we found that alterations in the directed functional connectivity between brain areas in the theta and alpha frequency bands are a core component of brain development at early stages of ASD (Sperdin et al., 2018). Higher activity within key nodes of the *social brain* (Adolphs, 2009; Brothers, 1990) for some toddlers and preschoolers with ASD was related to better visual exploration, and thus may represent a compensatory mechanism for ASD at such a young age (Sperdin et al., 2018; Welsh & Estes, 2018). Here, we used a data-driven, reference free EEG microstate approach (Michel & Koenig, 2018) to examine differences in the spatial organization and temporal dynamics of whole-brain neuronal networks in a large sample of toddlers and preschoolers with ASD and age-matched typically developing (TD) peers (N = 90, 3 years of age on average). EEG microstates represent the subsecond coherent activation within global functional brain networks and are usually defined in the literature as short-lasting periods (approx. 100 ms) of quasi-stable topographies of the electric potentials in the ongoing EEG (Wackermann et al., 1993; Lehmann et al., 2009). Interestingly, these rapidly changing EEG microstates are closely related and described as the electrophysiological correlates of fMRI resting-state networks (Britz et al., 2010; Musso et al., 2010; Bréchet et al., 2019). It is a commonly used method in the EEG field to study variations in the spatial organization and temporal dynamics of large-scale brain networks at rest or during a task and has provided important insights on how differently the brain processes information in populations with brain disorders (Michel & Koenig, 2018; Khanna et al., 2015). For example, numerous studies indicate that changes in the spatial and/or temporal characteristics of specific microstates represent critical markers for several brain disorders indicating that these spatial and temporal modulations may mirror how divergently a given individual with a neurodevelopmental condition is processing information compared to a TD individual (Khanna et al., 2015; Tomescu et al., 2015; Lehmann et al., 2005; Strelets et al., 2003; Stevens et al., 1997; Tomescu et al., 2014).

Building up on our previous study where we found alterations in the directed functional connectivity between brain areas in the theta and alpha frequency bands (Sperdin et al., 2018), we hypothesized that the toddlers and preschoolers with ASD would also show differences in the spatio-temporal properties of some microstates compared to their typically developing peers. We also looked at relationships between the temporal characteristics of the microstates and clinical phenotype. Finally, we used a bootstrapping approach (Schaer et al., 2015) to examine the stability of our findings. Post-hoc power analyses depending on the observed effect sizes were made to estimate the relationship between the sample size of our group and the observed statistical power. The bootstrapping procedure served to estimate the likelihood of finding the true result we observed in our full cohort from smaller sample sizes of participants.

## RESULTS

### Microstates analysis

The k-means cluster analysis across all participants identified five dominant maps, which explained 78.5 percent of the total variance (Figure 1). These five cluster maps correspond to the canonical microstate classes previously reported in the literature and were labeled accordingly (map A, B, C, D and E) (Michel & Koenig, 2018). These five cluster maps were used for further analysis.

**Figure 1.**
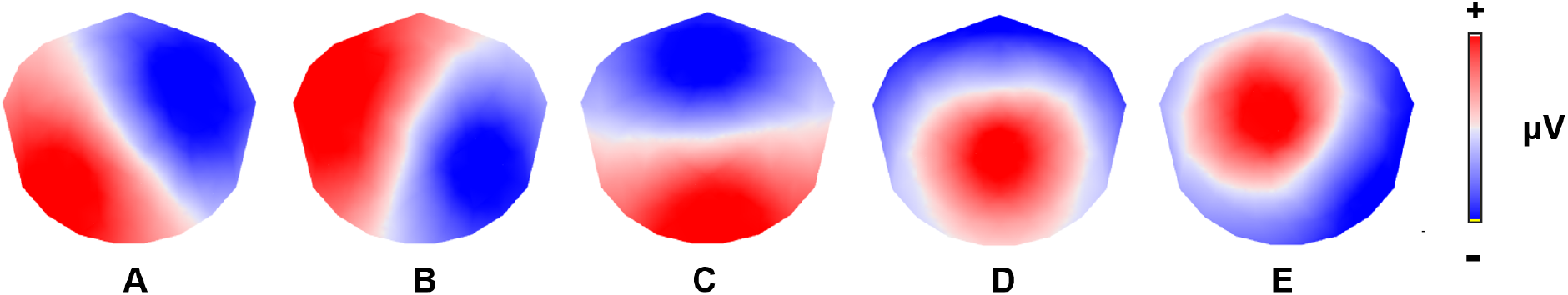
The five microstate topographies identified in the global clustering across all subjects.

When looking for group differences regarding the sum of temporal parameters, we found no difference between toddlers and preschoolers with ASD and their TD peers for the summed global explained variance (GEV) (df = 88, t = 1.633, p = 0.106) and the summed mean duration of all microstates (df = 88, t = 0.084, p = 0.933). However, we found a difference for the summed time coverage (df = 88, t = 3.305, p = 0.001) and the summed occurrence (df = 88, t = 2.904, p = 0.005) between both groups (Figure 2). The summed time coverage and the summed occurrence were lower in the toddlers and preschoolers with ASD compared to their TD peers. There was no significant difference for the time coverage when looking at the five maps separately between both groups (A : df = 88, t = 1.478, p = 0.143; B : df = 88, t = 1.936, p = 0.056; C : df = 88, t = 2.102, p = 0.038; D : df = 88, t = 0.803, p = 0.424; E : df = 88, t = 0.035, p = 0.973). The difference for the summed occurrences was driven by a significant difference in the occurrence of map C (df = 88, t = 2.821, p = 0.006) (Figure 3). Map C occurred significantly less in the toddlers and preschoolers with ASD compared to their TD peers. When considered separately, there was no significant differences in the frequency of occurrence for map A, B, D and E between both groups (A : df = 88, t = 1.829, p = 0.071; B : df = 88, t = 2.359, p = 0.021; D : df = 88, t = 1.207, p = 0.231; E : df = 88, t = 0.176, p = 0.861).

**Figure 2.**
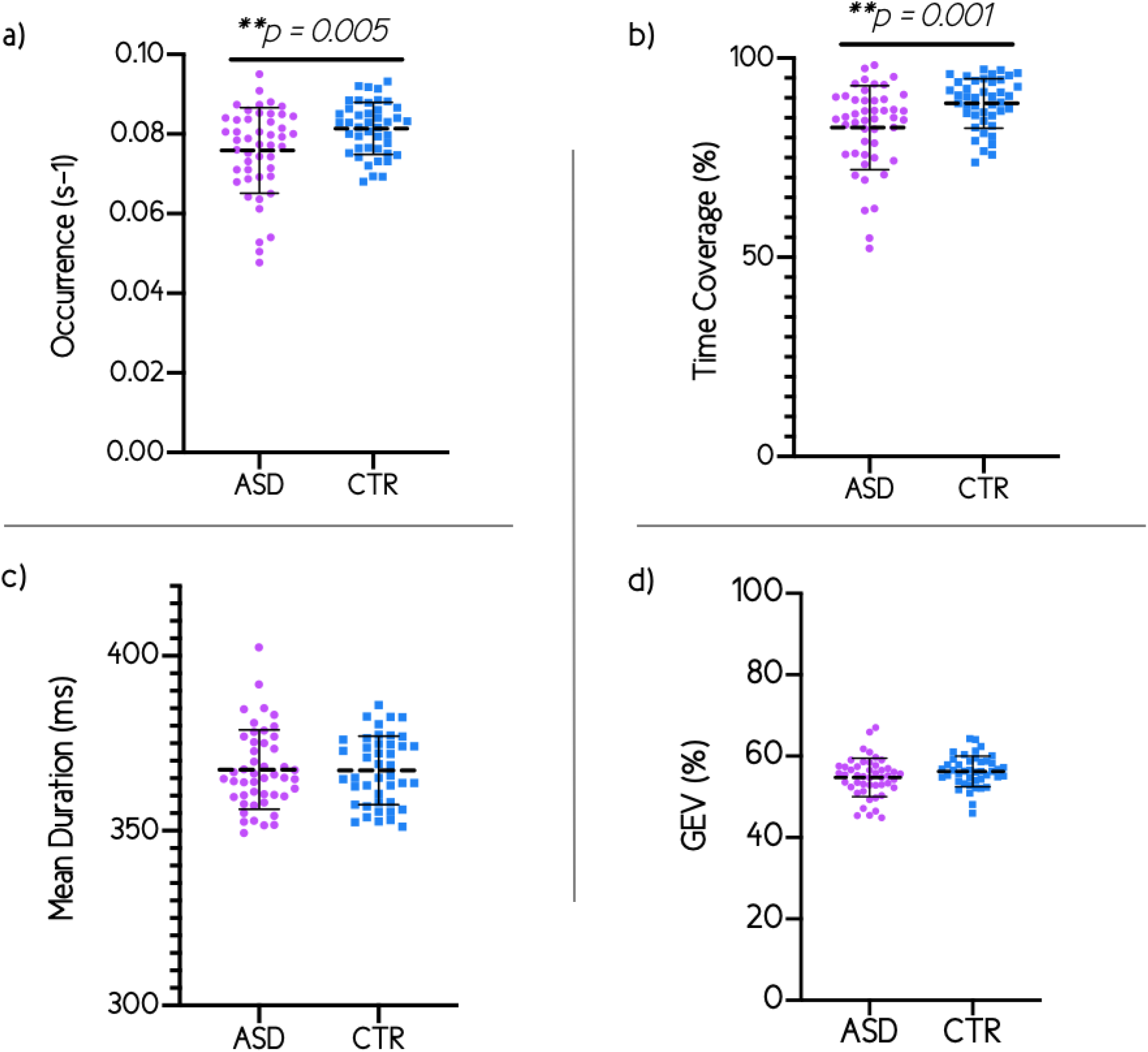
Results for the summed parameters a) Occurrence, b) Time coverage, c) Mean duration, d) Global explained variance.

**Figure 3.**
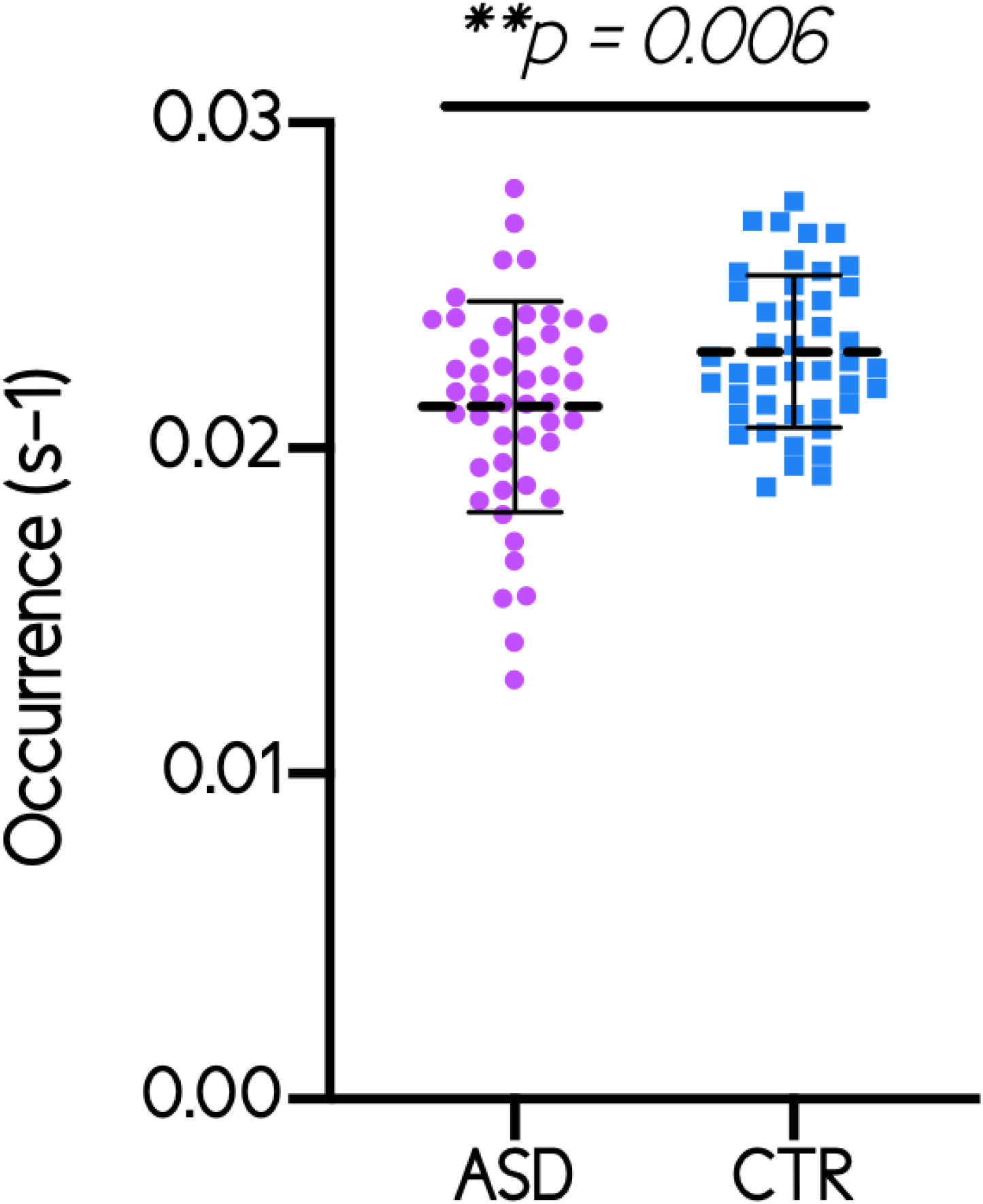
Difference of the occurrence of microstate class C between ASD toddlers and preschoolers and their TD peers.

### Correlation with clinical measures

The results of Spearman’s rank correlation revealed a positive association of the autism symptoms severity with map C. The occurrence of microstate class C significantly correlated with the ADOS total severity score (r = -0.359, N = 47, p = 0.013) (Figure 4). We also found a significant positive association between adaptive functioning level of toddlers and preschoolers with ASD and the occurrence of map C. The occurrence of microstate class C significantly correlated with the daily living skills subdomain of the Vineland Adaptive Behavior Scale-II (VABS-II) (r = 0.396, N = 47, p = 0.006) (Figure 5).

**Figure 4.**
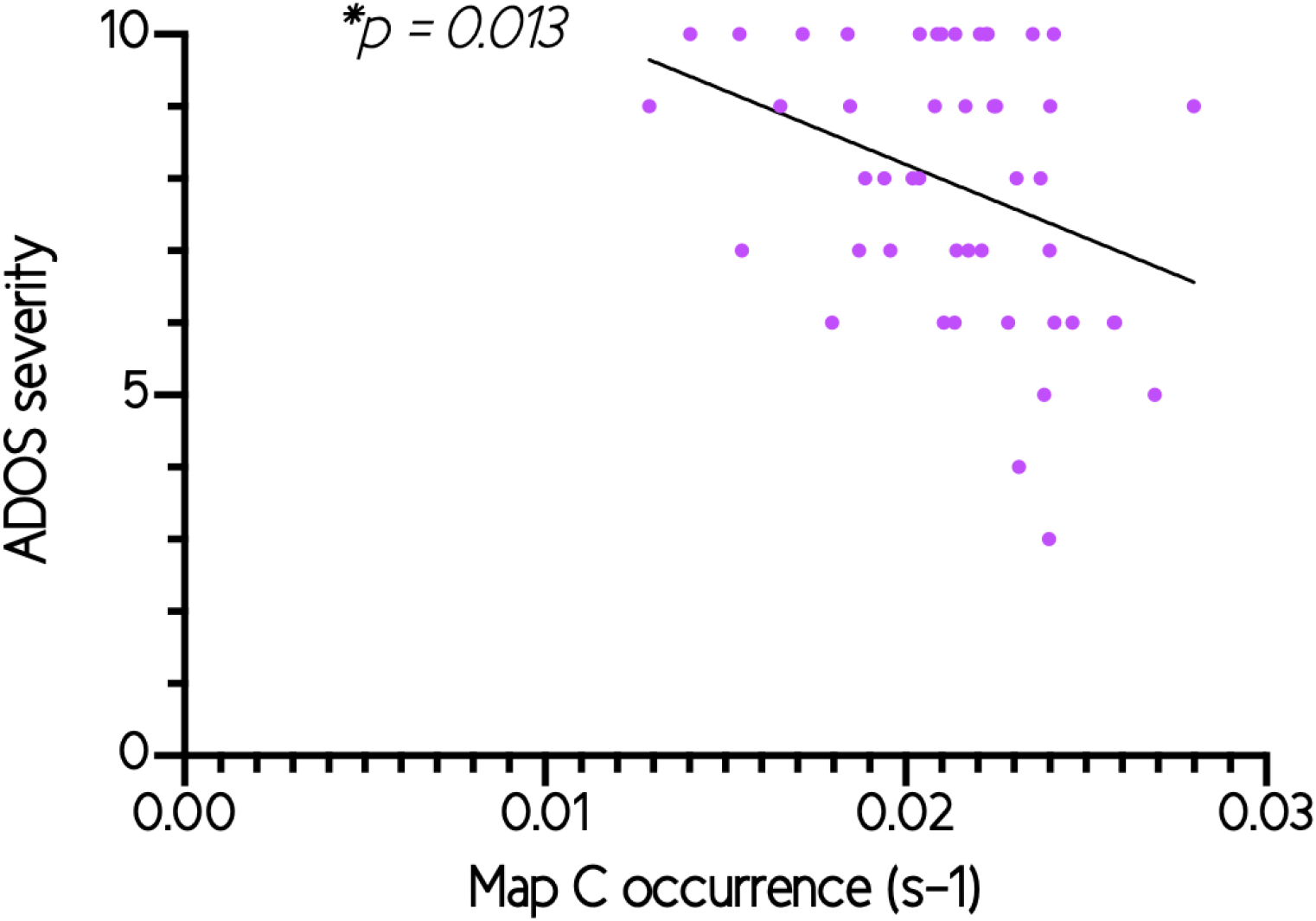
Negative correlation between the occurrence of microstate class C and the ADOS total severity score among toddlers and preschoolers with ASD.

**Figure 5.**
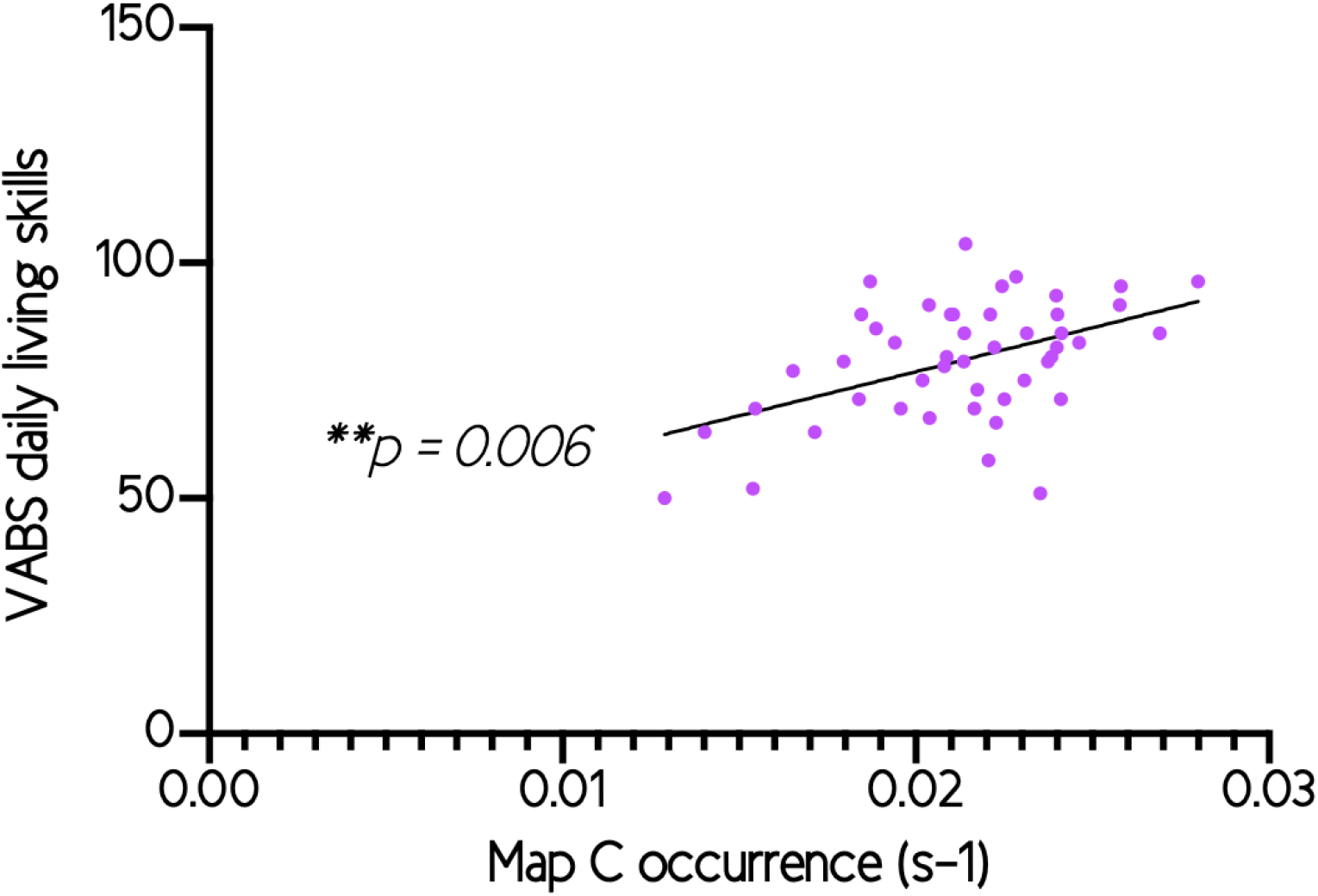
Positive correlation between the occurrence of microstate class C and the daily living skills VABS sub-domain among toddlers and preschoolers with ASD.

No association was found between the occurrence of map C and the developmental level of toddlers and preschoolers with ASD, using developmental quotient of the Mullen Scales of Early Learning composite score (MSEL) (r = 0.264, N = 39, p = 0.105). No other significant brain-behavioral relationship was found regarding the other maps (A, B, D and E).

### Bootstrapping analysis

The bootstrapping sub-sampling analysis tested the likelihood for observing a significant difference in the temporal parameters between both group for each simulated sample size (Figure 6). The results of stability analyses demonstrated a decrease in the likelihood of observing significant difference of temporal parameters between toddlers and preschoolers with ASD and TD peers as the sample size decreased. For instance, with a sample of 24 children with ASD and 24 TD children, a significant difference of Map C occurrence was only detected in 50 percent of the simulated sub-samples.

**Figure 6.**
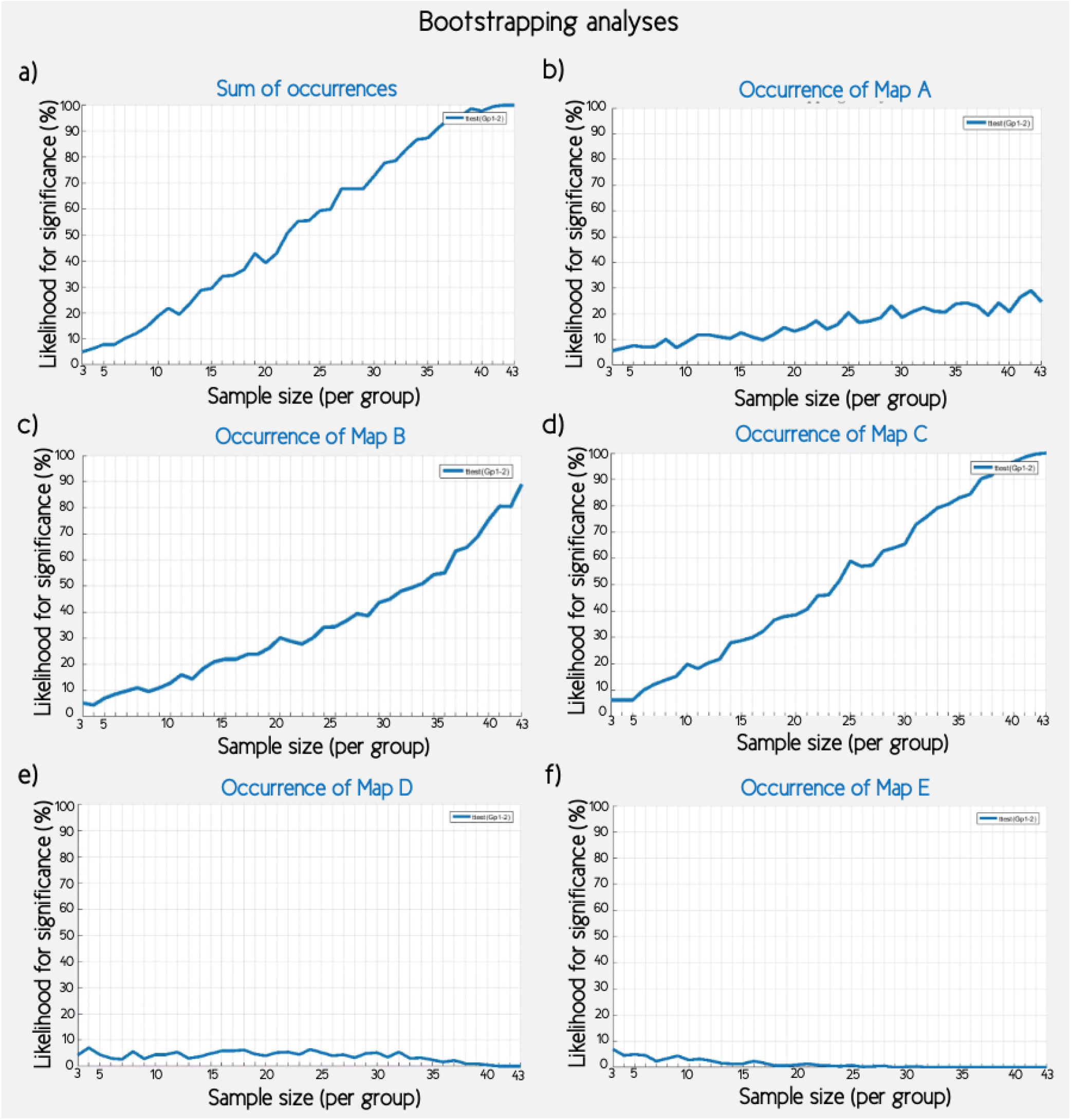
Bootstrapping analyses. The likelihood to observe a significant difference between toddlers and preschoolers with ASD and their TD peers, simulating sample sizes ranging from 3 to 43 individuals in each group, for parameters : a) Sum of occurrences, b) Occurrence of microstate class A, c) Occurrence of microstate class B, d) Occurrence of microstate class C, e) Occurrence of microstate class D, f) Occurrence of microstate class E.

## DISCUSSION

We applied a microstates analysis on EEG resting-state recordings acquired in toddlers and preschoolers with ASD and their TD peers (N = 90) and investigated modulations in four temporal parameters (the Global Explained Variance (GEV), the mean duration, the frequency of occurrence and the time coverage). The meta-criterion determined an optimal number of five template maps that best described the entire dataset explaining 78.5 percent of the global variance. The first four maps were identical in their spatial orientation to the canonical microstates classes A, B, C and D previously reported in the literature (Michel & Koenig, 2018; Khanna et al., 2015). The fifth map corresponds with microstate class E previously reported elsewhere (Bréchet et al., 2019; Custo et al., 2017). We found significant differences in the temporal parameters concerning the sum of occurrences of all maps and the sum of time coverage of all maps between toddlers and preschoolers with ASD and their TD peers. We didn’t find any difference regarding the sum of the GEV and the mean duration between any microstate class. In order to investigate the switch between maps and the dynamic of the brain in more detail, we focused on the frequency of occurrence of all the maps separately. We found that the difference between the groups in the summed occurrences was mostly driven by a difference in the occurrence of microstate class C. We didn’t find any significant difference regarding the frequency of occurrence for map A, B, D and E between toddlers and preschoolers with ASD and their TD peers.

Moreover, in the toddlers and preschoolers with ASD, we observed a negative relationship between the occurrence of microstate class C and the severity of ASD symptoms. We also found a positive relationship with the daily living skills suggesting that this microstate C occurred less in the toddlers and preschoolers with the most severe symptoms and adaptive behavior impairments. No other association was found between the occurrence of microstate C and the developmental level of toddlers and preschoolers with ASD. No other significant brain-behavioral relationship was found regarding the temporal parameters of the other microstate classes (A, B, D and E) and clinical, behavioural or developmental measures.

Recently, we published an exploratory study that combined eye-tracking and microstate analysis in a small sample of children (N = 28) (Jan et al., 2019). We found four group cluster maps, very similar to microstates classes A, B, C and D previously described in the literature (Jan et al., 2019). To the best of our knowledge, only three other studies using a microstate approach in individuals with ASD have been published (D’Croz-Baron et al., 2019; Jia & Yu, 2019; Malaia et al., 2016). D’Croz-Baron and colleagues 2019 found six microstate classes that best described their dataset in young adults with ASD and their TD peers. They found an increased occurrence for microstate classes B and E in individuals with ASD compared to their TD peers. There was a trend for microstate C being more present in their control group. However, this study only included 23 participants and only included adults. Our bootstrapping analysis demonstrated that a high level of likelihood to hit a significant result was reached with samples above approximately 30 participants per group. This would suggest that our sample size was sufficient in order to highlight significant differences between our groups of toddlers and preschoolers (N = 90).

In our study, microstate class C occurred less in the toddlers and preschoolers with ASD compared to their TD peers. In a seminal EEG-MRI study, Britz and colleagues 2010 linked microstate class C with positive BOLD activation in the anterior cingulate cortex, bilateral inferior frontal gyri and the insula (Britz et al., 2010). In the literature, the fronto-insular cortex has been related to the salience network (SN) (Seeley et al., 2007). More recently, microstate class C has been related to the Default Mode Network (DMN) (Bréchet et al., 2019; Custo et al., 2017; Seitzman et al., 2017), with bilateral activity in the lateral part of the parietal lobe and middle temporal gyrus found in EEG-fMRI study of Bréchet and colleagues 2019. The DMN is thought to play a role in “self-referential processing” (Padmanabhan et al., 2017) and is closely related to the SN and the central executive functions network (CEN) (Menon, 2011). The SN is suggested to be involved in switching between the CEN and the DMN in response to cognitive demands and therefore helps to direct attention towards salient (i.e.-behaviourally-relevant) stimuli (Sridharan et al., 2008; Menon & Uddin, 2010; Menon, 2011). Alterations in the SN and DMN have been described by functional and structural MRI studies in children, adolescents and adults with ASD (Uddin et al., 2017; Hull et al., 2017; Chen et al., 2017; Padmanabhan et al., 2017). Uddin and colleagues (2013) have suggested that functional hyperconnectivity within the SN and DMN networks as a distinguishing feature in children with ASD in comparison with their TD peers (Uddin et al., 2013). Taken together, there is an increasing number of evidence showing a different development of the SN and the DMN in individuals with ASD that may play a role in social deficits in ASD (Uddin et al., 2017; Padmanabhan et al., 2017).

Resting-state brain networks (RSNs) are usually studied using the functional MRI (fMRI) because of its high spatial resolution (Lee et al., 2013). However, these RSNs are sensitive to dynamic fluctuation (Preti et al., 2017) which are difficult to capture with fMRI because of moderate temporal resolution (in the order of seconds) and the delayed hemodynamic response. As such, EEG appears to be a valuable alternative technique to study RSNs because of its sub-second scale resolution (Abreu et al., 2020; Michel & Koenig, 2018). Here, we demonstrated a decreased occurrence of microstate class C among toddlers and preschoolers with ASD compared to their TD peers. This result suggests that the temporal properties of some specific microstate classes are altered in ASD at early developmental stages. Moreover, brain-behavioural analysis indicated that microstate class C, thought to reflect the DMN (Bréchet et al., 2019), occurred more in the toddlers and preschoolers with ASD who showed less symptoms and who had better daily living skill. Interestingly, in our directed functional connectivity study (Sperdin et al., 2018), we also found that young children with ASD who had less severe symptoms and a better visual exploration of social stimuli had more driving (i.e. hyper-connectivity) compared to the more affected ones between brain areas of the so-called *social brain* (Brothers, 1990; Adolphs, 2009). We suggested that the overall hyper-driving from certain brain regions might be a mechanism to compensate for the atypical development of the brain’s circuitry over time. The results presented here complement our initial findings and suggest that the less severely affected toddlers and preschoolers with ASD may also be able to compensate that way as more occurrence of microstate C was associated with less symptoms and better adaptive skills.

Highlighting brain differences early in life is important as it may ultimately help us to understand more what causes ASD and how the symptoms evolve over time given the vast heterogeneity of the ASD phenotype. We have started to characterize behavioural phenotypes in young children with ASD by taking developmental changes into account (Kojovic et al., 2020). Currently, we are exploring how these brain-network differences evolve over the course of development as we want to find out how the brains of very young children with ASD can compensate and how these mechanisms emerge. This will ultimately lead to the development of more individualized and thus adapted therapies early in life when the brain is most plastic.

## METHOD

### Participants

This study was approved by the Local Research Committee, the Commission Centrale d’Ethique de Recherche (CCER) in Geneva, Switzerland, and written informed consent was obtained from all children’s parents prior to inclusion in the study. 249 participants were recruited for the experiment. We didn’t manage to put the cap on 99 participants. We managed to put the cap on 150 participants (91 ASD and 59 TD). Out of those, we excluded 60 participants (44 ASD and 16 TD) because of either too many movement-related artefacts, noisy signal, lack of interest or insufficient amounts of epochs available for subsequent analysis. This was to be expected given the sensory processing issues frequently reported in this population (Kojovic et al., 2019). As a result, 90 participants were included in the final sample : 47 toddlers and preschoolers with ASD (11 females; mean age 2.92 years ±0.84, range 1.67-4.83) and 43 TD peers (15 females; mean age 3.00 years ±1.31, range 1.08-5.58). Groups did not differ by age (p = 0.723) and gender (p = 0.250). Five minutes of spontaneous EEG recordings were acquired for all the participants included in the study. All participants were recruited as a part of the Geneva Autism Cohort, a longitudinal cohort of young children (Robain et al., 2020; Franchini et al., 2016). Toddlers and preschoolers were included in the ASD group if the previously established clinical diagnosis was confirmed by exceeding the threshold limit for ASD on ADOS-G (Autism Diagnostic Observation Schedule-Generic) (Lord et al., 2000) or ADOS-2 (Second version) (Lord et al., 2012). The ADOS assessments were performed and scored by experienced clinicians working in the research team and specialized in ASD identification. For toddlers and preschoolers who were administered the ADOS-G assessment, the scores were recoded according to the revised ADOS algorithm (Gotham et al., 2007, 2009) to ensure comparability with ADOS-2. The mean Severity Score at ADOS for the toddlers and preschoolers with ASD group was 7.94 ± 1.86. For the control group, TD toddlers and preschoolers were recruited through announcements in the Geneva community. They were also assessed by ADOS-G or ADOS-2, to ensure the absence of ASD symptoms, which would be an exclusion criterion. All TD participants had a minimal severity score of 1, except one child who had a score of 2. Children were excluded from the control group if they presented any neurological/psychiatric conditions and learning disabilities according to parents’ interview and questionnaire, or if they had a sibling or first-degree parent diagnosed with ASD. The assessment of all participants also included adaptive behavior using the Vineland Adaptive Behaviour Scale-II (VABS-II) (Sparrow, 2011). The VABS-II is a standardized parent report interview which measures adaptive behavior level in socialization, communication, daily living skills and motor sub-domains. Finally, to estimate developmental level, all participants were assessed with the Mullen Scales of Early Learning (MSEL) (Lee, 2013). The MSEL is a measure of cognitive functioning for children from birth through age 68 months, including gross-motor, visual reception, fine motor, receptive language, and expressive language scales sub-domains. See Table 1 for characteristics of study participants.

**Table 1.**
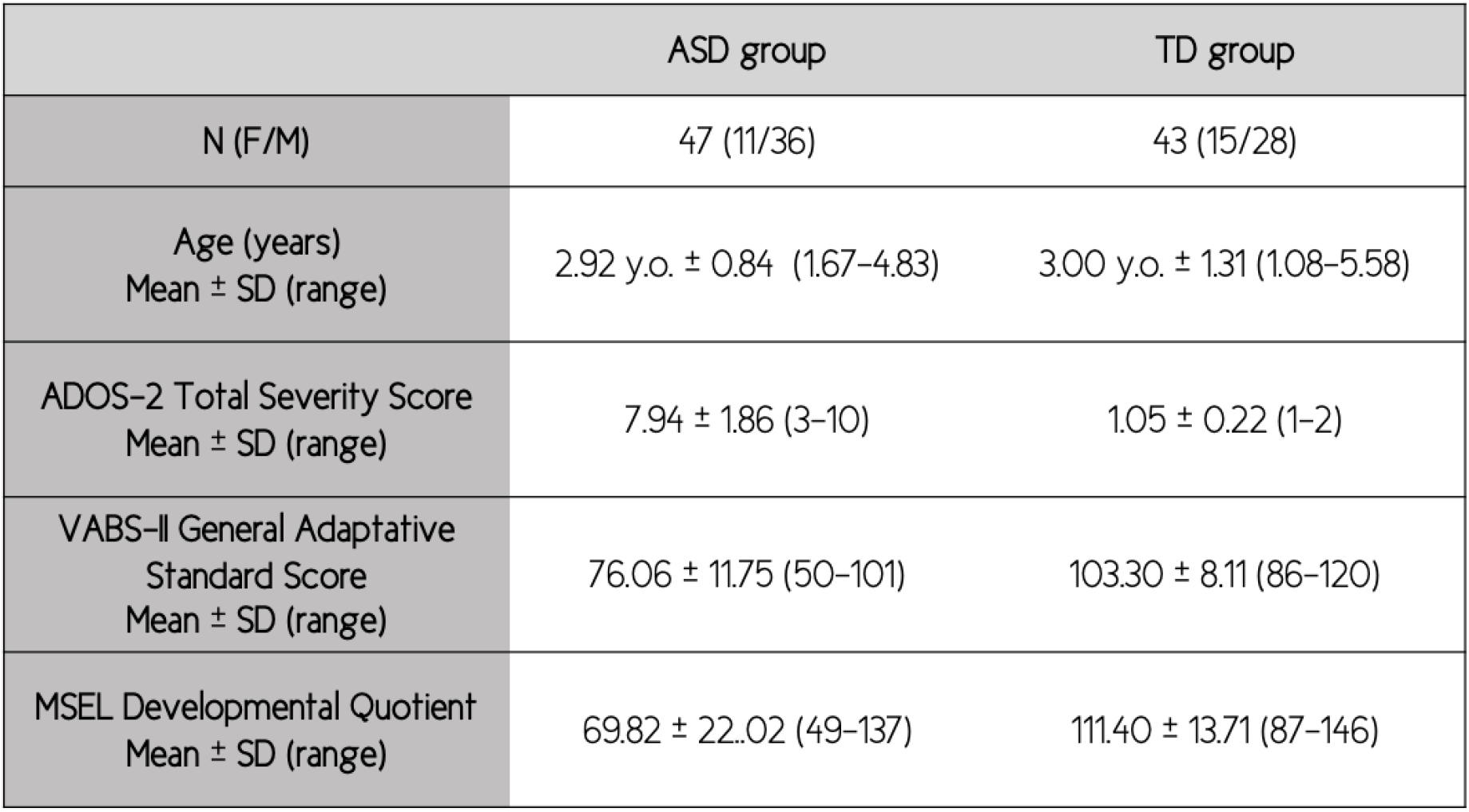
Characteristics of study participants.

|  | ASD group | TD group |
| --- | --- | --- |
| N (F/M) | 47 (11/36) | 43 (15/28) |
| Age (years)<br>Mean ± SD (range) | 2.92 y.o. ± 0.84 (1.67–4.83) | 3.00 y.o. ± 1.31 (1.08–5.58) |
| ADOS-2 Total Severity Score<br>Mean ± SD (range) | 7.94 ± 1.86 (3–10) | 1.05 ± 0.22 (1–2) |
| VABS-II General Adaptive<br>Standard Score<br>Mean ± SD (range) | 76.06 ± 11.75 (50–101) | 103.30 ± 8.11 (86–120) |
| MSEL Developmental Quotient<br>Mean ± SD (range) | 69.82 ± 22.02 (49–137) | 111.40 ± 13.71 (87–146) |

### Procedure and task

The experiment was conducted in a quiet room. To help the children and their relatives to get familiar with the protocol, they received two weeks prior to their visit a kit containing a custom handmade EEG cap, pictures and a video illustrating the experiment. Participants were seated alone on a comfortable seat or on their parents lap in order to reassure them and keep them as calm as possible to avoid hand and body movements. Once seated, the experimenter measured the circumference of the head. The cap of the corresponding size was then prepared and gently placed on the participant’s head. A couple of minutes were taken in order to allow the participants to settle into the experiment’s environment and get used to the cap before starting the experiment. To best capture the child’s attention during the experiment, we showed them an age-appropriate animated cartoon of their choice. Impedance were measured and electrodes adjusted to keep values below 50 k*Ohm*.

### EEG acquisition and preprocessing

The EEG was acquired with a Hydrocel Geodesic Sensor Net (HCGSN, Electrical Geodesics, USA) with 129 scalp electrodes at a sampling frequency of 1000*Hz*. On-line recording was band-pass filtered at 0–100*Hz* using the vertex as reference. Data pre-processing and microstate analysis were done using Cartool (http://sites.google.com/site/cartoolcommunity/) and Matlab (Natick, MA). First, we down-sampled the montage to a 110-channel electrode array to exclude electrodes on the cheek and the neck since those are often contaminated with muscular artefacts. Data were filtered between 1 and 40*Hz* (using Butterworth filters) and a 50*Hz* notch filter was applied. Each file was then visually inspected to detect periods of movement artefacts.These periods were excluded. We performed Independent component analysis (ICA) on the data to identify and remove the components related to eye movement artefacts (eye blinks, saccades) (Jung et al., 2000, 1996). Channels with substantial noise were interpolated using spherical spline interpolation for each recording. The cleaned data were down-sampled to 125*Hz*, recalculated against the average reference and a spatial filter was applied. Finally, EEG experts (HFS, AB) reinspected all data to ensure that no artefacts had been missed.

### Microstates analysis

The pipeline for the analysis is illustrated in (Figure 7). We applied a k-means cluster analysis to the data of each subject to estimate the optimal set of topographies explaining the EEG signal. The clustering was applied only at local maxima of the Global Field Power (GFP) which is calculated as the standard deviation of all electrodes at a given time point and represents time points of highest signal-to-noise ratio (see (Murray et al., 2008) for formulas). The polarity of the maps was ignored during the procedure. The k-means cluster analysis was first computed at the individual level and then across all participants (children with ASD and TD children together) to obtain the group cluster maps. In order to determine the optimal number of maps at both levels (both within and across subjects), we applied a meta-criterion that includes seven independent criteria. For a detailed description of these criteria, see (Custo et al., 2017) and (Bréchet et al., 2019). Then, the cluster maps for all participants were fitted back to the original EEG of each subject. This way that spatial correlation was calculated between the cluster maps and each individual data point and that data point was labeled with the cluster map that showed the highest correlation. All data points were included at the exception of periods marked as artefacts during the preprocessing. Polarity of the maps was again ignored for the back-fitting procedure. Data points that didn’t correlate more than 50 percent with a given group cluster map were marked as unlabeled.

**Figure 7.**
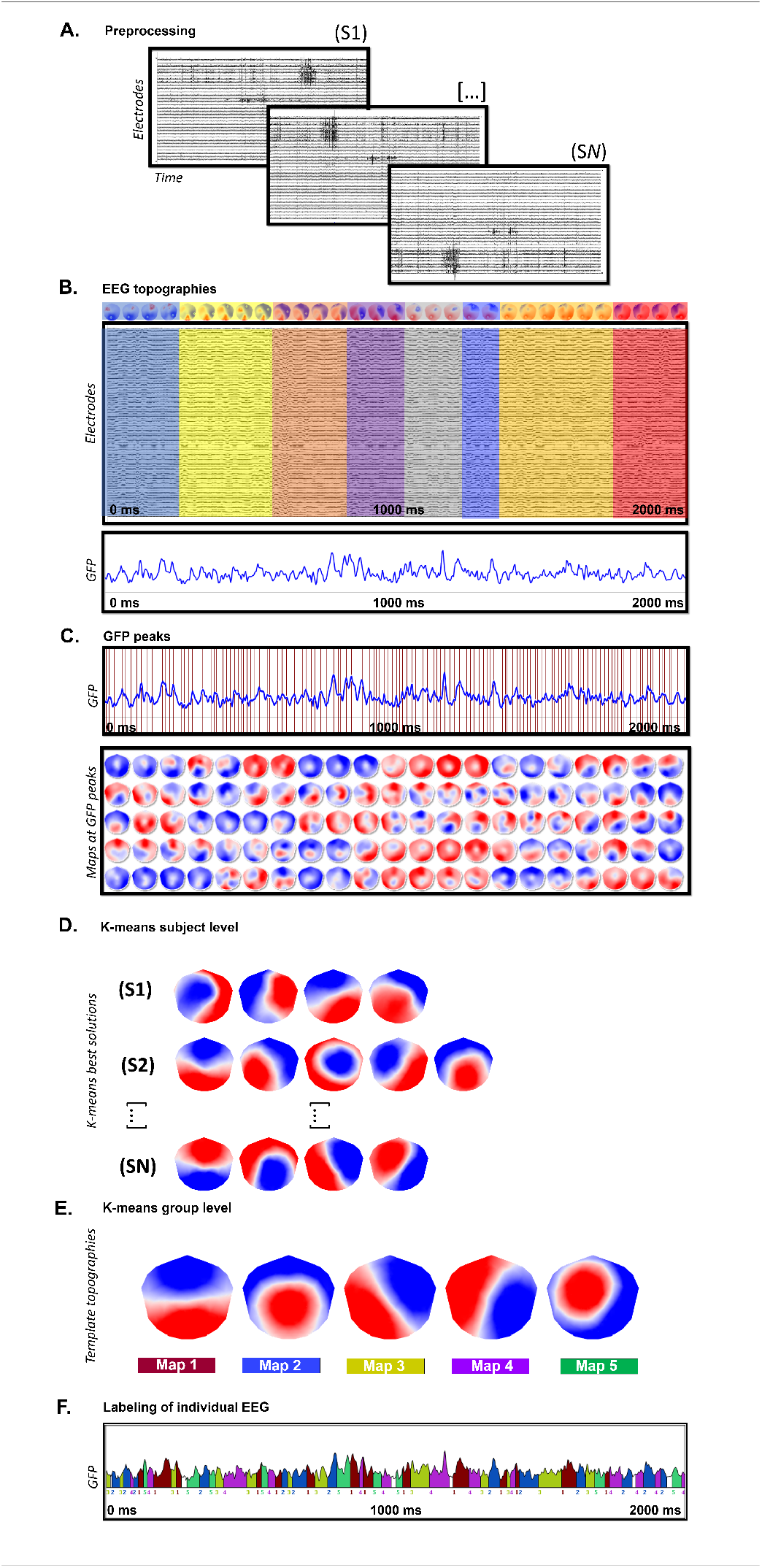
Microstate analysis: (A) Standard preprocessing of all the acquired high-density EEGs (110 channels). (B) A 2-sec cleaned EEG and its corresponding global field power (GFP). Periods of quasi stable map topographies (on top) are superimposed on the cleaned EEG and marked in different colors. (C) For each individual recording, peaks of GFP were determined (red vertical lines) and their specific potential maps were selected and submitted to a k-means clustering procedure (D). The best k-means clustering solutions at the individual level were selected based on the meta-criterion. (E) The best solutions obtained for each subject in step (D) are submitted altogether to a second k-means group cluster analysis. The meta-criterion identified a best solution with 5 template topographies (microstate classes). (F) The template topographies obtained in (E) are fitted back to the individual EEG recording and each time point is labeled with the cluster map having the highest spatial correlation (winner-takes-all). The microstate sequence is used, for every subject, to extract the temporal parameters and statistical analysis.

Four temporal parameters of the microstates were computed for each individual recording : GEV, the mean duration, the time coverage and the frequency of occurrence. The GEV is an estimate of the explained variance of a given map, weighted by the GFP. The mean duration is the average duration in milliseconds that a given cluster map is continuously present. The time coverage is the percentage of total time for a given cluster map in the individual EEG recording. The frequency of occurrence represents the number of times per second that a given cluster map appears in the individual EEG recording (Michel & Koenig, 2018).

### Statistical analysis

We first checked if the temporal parameters of the microstate classes had a normal distribution using with Kolmogorov-Smirnov tests. We summed each of the four temporal parameters separately to obtain the sum of the GEV, the mean duration, the time coverage and occurrence of all maps. We investigated group differences using unpaired t-tests for the sum of each temporal parameters applying Bonferoni’s multiple correction to take into account the four parameters (p-value significant if < 0.0125). We then observed the group difference for the occurrence parameter for each map separately using unpaired t-tests and applying Bonferoni’s multiple correction to take in account the five maps (p-value significant if < 0.01).

We investigated possible brain-behavioral relationships between the temporal parameters of the microstate classes with autism symptom severity scores. Among toddlers and preschoolers with ASD, we correlated autism symptom severity, adaptive functioning level and developmental level with the temporal parameters which were statistically different between ASD and TD groups. As the distribution of clinical data were not normal, we applied two-tailed Spearman’s rank correlation between the occurrence of microstates and Autism Diagnostic Observation Schedule total severity score (ADOS-G and ADOS-2) (Lord et al., 2000, 2012)), daily living skills sub-domain of the Vineland Adaptive Behavior Scales (VABS-II) (Sparrow, 2011) and developmental quotient of the Mullen Scales of Early Learning composite score (MSEL) (Lee, 2013).

Finally, given the large heterogeneity in the ASD phenotype, we wanted to estimate the likelihood of finding the significant results we observed in our full cohort from smaller sample sizes of participants. To do so, we simulated sample sizes ranging from 3 to 43 individuals in each group (with steps of 1 participant), using 500 bootstrapped sub-samples for each sample size. With each sample, unpaired t-test were performed to assess the significance of difference between both group, using a statistical threshold of p *<* 0.05.

## Acknowledgements

The authors would like thank all the families who took part in this research. This research was supported by the National Center for Competence in Research “Synapsy”, financed by the Swiss National Science Foundation (SNF, 51NF40-185897) and by private funding by the Fondation Pole Autisme (http://www.pole-autisme.ch). This work was further supported by individual SNF grants to Marie Schaer (#163859 & #190084) and to Aurélie Bochet (#323530-183979).

## Author contributions

Conception and design of the experiment: H.F.S, T.R and M.S.; Acquisition of data H.F.S, A.B, T.R.; Analysis and/or interpretation of data: A.B, H.F.S, M.S., C.M.; Drafting of the manuscript: A.B and H.F.S.; All authors revised the manuscript critically for important intellectual content.

